# Common genomic regions underlie natural variation in diverse toxin responses

**DOI:** 10.1101/325399

**Authors:** Kathryn S. Evans, Shannon C. Brady, Joshua S. Bloom, Robyn E. Tanny, Daniel E. Cook, Sarah E. Giuliani, Stephen W. Hippleheuser, Mostafa Zamanian, Erik C. Andersen

**Author notes:** Corresponding author: Erik C. Andersen Department of Molecular Biosciences Northwestern University 3125 Cook Hall 2205 Tech Drive Evanston, IL 60208. The first two authors contributed equally.

## Abstract

Phenotypic complexity is caused by the contributions of environmental factors and multiple genetic loci, interacting or acting independently. Studies of yeast and *Arabidopsis* found that the majority of natural variation across phenotypes is attributable to independent additive quantitative trait loci (QTL). Detected loci in these organisms explain most of the estimated heritable variation. By contrast, many heritable components underlying phenotypic variation in metazoan models remain undetected. Before the relative impacts of additive and interactive variance components on metazoan phenotypic variation can be dissected, high replication and precise phenotypic measurements are required to obtain sufficient statistical power to detect loci contributing to this missing heritability. Here, we used a panel of 296 recombinant inbred advanced intercross lines of *Caenorhabditis elegans* and a high-throughput fitness assay to detect loci underlying responses to 16 different toxins, including heavy metals, chemotherapeutic drugs, pesticides, and neuropharmaceuticals. Using linkage mapping, we identified 82 QTL that underlie variation in responses to these toxins and predicted the relative contributions of additive loci and genetic interactions across various growth parameters. Additionally, we identified three genomic regions that impact responses to multiple classes of toxins. These QTL hotspots could represent common factors impacting toxin responses. We went further to generate near-isogenic lines and chromosome-substitution strains and then experimentally validated these QTL hotspots, implicating additive and interactive loci that underlie toxin-response variation.

## INTRODUCTION

Rapid advances in sequencing technologies enabled the collection of high-quality genomic datasets for many species (Mardis 2017). These data, paired with a broad range of high-throughput phenotypic assays, made quantitative genetics a powerful tool in biology. Linkage mapping has been used to identify quantitative trait loci (QTL), leading to profound impacts on human health (Easton *et al.* 1993; Cowley 2006; Altshuler, Daly, and Lander 2008), agriculture and livestock (Rothschild, Hu, and Jiang 2007; Johnsson *et al.* 2015; Leal-Bertioli *et al.* 2015; Shang *et al.* 2016), and basic biology (T. F. Mackay 2001; Peng *et al.* 2016; Andersen *et al.* 2015). Despite the growing number of detected QTL across numerous traits, these QTL often do not explain the complete heritable component of trait variation (Rockman 2012). This missing heritability can be attributed to undetected small-effect additive loci and/or interactions between QTL (Bloom *et al.* 2015). Although some studies contend that epistatic effects among QTL might explain missing heritability (Nelson, Pettersson, and Carlborg 2013; T. F. C. Mackay 2015; Lachowiec *et al.* 2015; Malmberg *et al.* 2005; Zuk *et al.* 2012), others argue that missing heritability comprises small-effect additive loci that remain undetected in cases where statistical power is too low (Hill, Goddard, and Visscher 2008; Mäki-Tanila and Hill 2014; Ehrenreich 2017; Yang *et al.* 2010). Quantitative geneticists have leveraged large numbers of recombinant strains in both yeast and *Arabidopsis* to overcome power limitations and concluded that, when power is sufficient, small-effect additive components can be identified that account for nearly all of the heritability of a given trait (Bloom *et al.* 2015, 2013; Simon *et al.* 2008). We require a metazoan system with high statistical power to determine whether this predominantly additive-QTL model remains broadly applicable in animals.

One such tractable metazoan is the roundworm nematode *Caenorhabditis elegans.* The genetic variation among a panel of recombinant inbred advanced intercross lines (RIAILs) generated between the N2 and CB4856 strains of *C. elegans* (Rockman and Kruglyak 2009; Andersen *et al.* 2015) has been leveraged in many linkage mapping analyses (McGrath *et al.* 2009; Bendesky *et al.* 2011; Lee *et al.* 2017; Bendesky *et al.* 2012; K. D. Singh *et al.* 2016; Schmid *et al.* 2015; Balla *et al.* 2015; Zdraljevic *et al.* 2017; Zamanian *et al.* 2018a; Andersen *et al.* 2014; Bendesky and Bargmann 2011; Viñuela *et al.* 2010; Doroszuk *et al.* 2009; Snoek *et al.* 2014; Rodriguez *et al.* 2012; Kammenga *et al.* 2007; Gutteling, Riksen, *et al.* 2007; Li *et al.* 2006; Gutteling, Doroszuk, *et al.* 2007; Glater, Rockman, and Bargmann 2014; Reddy *et al.* 2009; Rockman, Skrovanek, and Kruglyak 2010; Seidel *et al.* 2011; Seidel, Rockman, and Kruglyak 2008). Additionally, a high-throughput phenotyping platform to rapidly and accurately measure animal fitness could provide the replication and precision required to detect small-effect additive loci and to determine the relative contributions of additive and/ or epistatic loci to trait variation (Andersen *et al.* 2014; Zdraljevic *et al.* 2017). Notably, the combination of this panel and phenotyping platform have facilitated linkage mappings of multiple distinct fitness parameters, resulting in the detection of a single QTL, in fact a single quantitative trait gene (QTG), that underlies several fitness-related traits (Andersen *et al.* 2014; Zdraljevic *et al.* 2017). This example of pleiotropy suggests that large-scale studies could reveal additional pleiotropic effects.

Such large-scale studies have implicated pleiotropic QTL that impact the expression of a broad range of genes (Keurentjes *et al.* 2007; Breitling *et al.* 2008; Rockman, Skrovanek, and Kruglyak 2010; Hasin-Brumshtein *et al.* 2016). Variation in the master regulators that are within these expression QTL hotspots have downstream effects on the transcription of many genes. Similarly, other QTL hotspots could impact multiple traits, such as responses to various conditions. In yeast, most chemical-response QTL are thought to be unique to one or a few conditions, whereas few QTL have been found to have pleiotropic effects across many conditions (Ehrenreich *et al.* 2012; U. M. Singh *et al.* 2017; Knoch *et al.* 2017). Although QTL underlying responses to individual conditions have been identified across multiple animal models (Najarro *et al.* 2015; Marriage *et al.* 2014; Highfill *et al.* 2017; Bubier *et al.* 2014; Crusio *et al.* 2016), the existence of QTL hotspots that influence multiple condition responses has yet to be observed broadly in metazoans.

Here, we performed a set of linkage-mapping experiments with a large panel of recombinant lines to identify QTL implicated in responses to 16 different toxins and found three QTL hotspots that underlie many of these responses. We demonstrated how high replication in the high-throughput fitness assay can enable the identification and validation of QTL, even in cases of small phenotypic effects. Additionally, we analyzed relative contributions of additive and epistatic genetic loci in various toxin responses. Finally, we discovered evidence for interactions between loci of the N2 and CB4856 strains that impact several toxin responses and could suggest how large regions of the genome were swept across the species.

## MATERIALS AND METHODS

### Strains

Animals were grown at 20°C using OP50 bacteria spotted on modified nematode growth medium (NGMA), containing 1% agar and 0.7% agarose to prevent animals from burrowing. For each assay, strains were propagated for five generations after starvation to reduce transgenerational effects of starvation (Andersen *et al.* 2014). Recombinant inbred advanced intercross lines (RIAILs) used for linkage mapping were constructed previously (Andersen *et al.* 2015). The construction of near-isogenic lines (NILs) and chromosome substitution strains (CSSs) is detailed below, and all strains are listed in the **Supplementary Information**. Strains are available upon request.

### High-throughput toxin response assay

We used a modified version of the high-throughput fitness assay described previously (Zdraljevic *et al.* 2017). Populations of each strain were passaged for four generations, amplified, and bleach-synchronized. Approximately 25 embryos from each strain were then aliquoted to 96-well microtiter plates at a final volume of 50 µL of K medium (Boyd, Smith, and Freedman 2012). Embryos hatched overnight and arrested in the L1 larval stage. The following day, arrested L1 animals were fed HB101 bacterial lysate (Pennsylvania State University Shared Fermentation Facility, State College, PA; (García-González *et al.* 2017)) at a final concentration of 5 mg/mL in K medium and were grown to the L4 larval stage for 48 hours at 20°C with constant shaking. Three L4 larvae were then sorted using a large-particle flow cytometer (COPAS BIOSORT, Union Biometrica, Holliston, MA) into microtiter plates that contained HB101 lysate at 10 mg/mL, K medium, 50 µM kanamycin, and either diluent (1% DMSO or 1% water) or diluent and a toxin of interest. The sorted animals were then grown for 96 hours at 20°C with constant shaking. During this time, the sorted animals matured to adulthood and laid embryos, yielding a population of parent and progeny in each microtiter well. Prior to the measurement of fitness parameters from the populations, animals were treated with sodium azide (50 mM in M9) to straighten their bodies for more accurate growth-response parameter measurements. Traits that were measured by the BIOSORT include brood size (n), animal length (time of flight, TOF), and optical density (extinction time, EXT).

### Toxin-response trait calculations

Phenotypic measurements collected by the BIOSORT were processed using the R package *easysorter*, which was specifically developed for processing this type of data set (Shimko and Andersen 2014). Briefly, the function *read_data* imported raw phenotypic data then identified and eliminated bubbles. Next, the *remove_contamination* function discarded wells that contained bacterial or fungal contamination (determined by visual inspection) prior to analyzing population parameters. The *sumplate* function then calculated normalized measurements and summary statistics of the assayed traits for the population of animals in each well. The number of animals in each well was divided by the number of animals sorted into that well, yielding a normalized brood size (norm.n). Additionally, optical density (EXT) of each animal was divided by animal length (TOF), resulting in a normalized optical density (norm.EXT) for each animal in each well. The norm.EXT measurement represents the optical density without conflating variation in body length. The summary statistics calculated for each population include 10th, 25th, 50th, 75th, and 90th quantiles, mean, and median measurements of TOF, EXT, and norm.EXT as well as variance for TOF and EXT. Previously, each of these summary statistics has been shown to reveal distinct genetic architectures underlying trait variation, suggesting values to demonstrate the range of biological phenomena that can be captured using this platform (Andersen *et al.* 2015). In total, this analysis resulted in 24 phenotypic measurements for each condition tested. When strains were measured across multiple assay days, the *regress(assay=TRUE)* function was used to fit a linear model with the formula (*phenotype ∼ assay*) to account for differences among assays. Next, outliers were removed by eliminating phenotypic values that were outside two standard deviations of the mean (unless at least 5% of the strains were outside this range in the case of RIAIL assays). Finally, toxin-specific effects were calculated using the *regress(assay=FALSE)* function from *easysorter*, which fits a linear model with the formula (*phenotype ∼ control phenotype*) to generate residual phenotypic values that account for differences between populations that were present in control conditions. For this reason, strain phenotypes in control conditions can influence regressed toxin effects and trait categorizations (below).

### Dose-response assays

For each toxin, a dose-response experiment was performed using quadruplicates of four genetically diverged strains (N2, CB4856, DL238, and JU258). Animals were assayed using the high-throughput fitness assay, and toxin-response trait calculations were performed as described above (**File S1**). The concentration of each toxin that provided a highly reproducible toxin-specific effect with variation between N2 and CB4856 across three distinct traits (brood size - norm.n, mean length - mean.TOF, and mean optical density - mean.norm.EXT) was selected for linkage mapping experiments. The chosen concentrations and diluents of each toxin are as follows: cadmium 100 µM in water, carmustine 250 µM in DMSO, chlorothalonil 250 µM in DMSO, chlorpyrifos 1 µM in DMSO, cisplatin 250 µM in water, copper 250 µM in water, diquat 250 µM in water, fluoxetine 250 µM in DMSO, FUdR 50 µM in water, irinotecan 125 µM in DMSO, mechlorethamine 200 µM in DMSO, paraquat 500 µM in water (50 µM was used for the CSS and NIL assays), silver 150 µM in water, topotecan 400 µM in water, tunicamycin 10 µM in DMSO, and vincristine 80 µM in water (**Table S1**). The concentration of paraquat differs the concentration used previously (Andersen *et al.* 2015), suggesting why the genetic architectures are different between the two studies. Toxins assayed in this manuscript were purchased from Fluka (chlorothalonil, #36791-250MG; chlorpyrifos, #45395-250MG; diquat dibromide monohydrate, #45422-250MG-R), Sigma (vincristine sulfate salt, #V8879-25MG; cisplatin, #479306-1G; silver nitrate, #209139; carmustine, #1096724-75MG; topotecan hydrochloride, #1672257-350MG), Calbiochem (tunicamycin, #654380), Aldrich (mechlorethamine hydrochloride, #122564-5G, cadmium chloride #01906BX), Alfa Aesar (irinotecan hydrochloride trihydrate, #AAJ62370-MD), Bioworld (5-fluoro-2’-deoxyuridine, #50256011), Enzo Life Sciences (fluoxetine, #89160-860), Mallinckrodt (cupric sulfate, #4844KBCK), and Chem Service (paraquat, #ps-366).

### Principal Component Analysis of RIAILs

A total of 296 RIAILs were assayed in the high-throughput fitness assay described previously in the presence of each toxin listed above as well as control conditions (water or DMSO, **File S2**). Because some of the 24 population parameters measured by the BIOSORT are highly correlated, a principal component analysis (PCA) was performed. For each growth-response trait, RIAIL phenotypic measurements were scaled to have a mean of 0 and a standard deviation of 1. The *princomp* function within the *stats* package in R (R Core Team 2017) was used to run a principal component analysis for each toxin. For each toxin, the minimum number of principal components (PCs) that explained at least 90% of the total phenotypic variance in the RIAILs was mapped through linkage mapping (**File S3**, **Table S2**). A total of 97 PCs were mapped.

### Linkage mapping

Linkage mapping was performed on each of the 97 PCs (described above) using the R package *linkagemapping* (www.github.com/AndersenLab/linkagemapping, **File S4**). The genotypic data and residual phenotypic data were merged using the *merge_pheno* function. Quantitative trait loci (QTL) were detected using the *fsearch* function, which scaled phenotypes to have a mean of zero and variance of one, then calculated logarithm of odds (LOD) scores for each marker and each trait as, where r is the Pearson correlation coefficient between RIAIL genotypes at the marker and trait values (Bloom *et al.* 2013). We note that this scaling of the data did not impact mappings in that scaled mappings and unscaled mappings were identical. The phenotypic values of each RIAIL were then permuted randomly while maintaining correlation structure among phenotypes 1000 times to calculate a significance threshold based on a genome-wide error rate of 5%. This threshold was set for each mapped PC independently to avoid biases introduced by performing large numbers of mappings. The marker with the highest LOD score was then set as a cofactor and mapping repeated iteratively until no significant QTL were detected. Finally, the *annotate_lods* function was used to calculate the fraction of variation in RIAIL phenotypes explained by each QTL. 95% confidence intervals were defined by markers within a 1.5-LOD drop from the marker with the maximum LOD score. We additionally performed a two-dimensional genome scan using the function *scantwo()* in the *qtl* package (Broman *et al.* 2003) for all 47 significantly mapped PCs (**File S5**). Significant interactions were determined by permuting the phenotype data for each PC 1000 times and determining the 5% genome-wide error rate for QTL detection.

### Heritability estimates

Broad-sense heritability was estimated for each of the 97 PCs using the formula H^2^ = (σ_R_^2^ -σ_P_^2^)/σ_R_^2^ where σ_R_^2^ and σ_P_^2^ are the variance among the RIAIL and parental (N2 and CB4856) phenotypic values, respectively (Brem and Kruglyak 2005). A variance component model using the R package *regress* was used to estimate the fraction of phenotypic variation explained by additive genetic factors (‘narrow-sense’ heritability) (David Clifford And 2014, 2006; Bloom *et al.* 2015). The additive relatedness matrix was calculated as the correlation of marker genotypes between each pair of strains. In addition, a two-component variance component model was calculated with both an additive and pairwise-interaction effect (**File S6**). The pairwise-interaction relatedness matrix was calculated as the Hadamard product of the additive relatedness matrix.

### Calculation of hotspots

We estimated cM distances from recombination events in the RIAIL panel to account for non-uniform distribution of genetic diversity across the genome. The genome was divided into 65 total bins with each bin containing 26 cM. To determine if the 82 QTL significantly clustered around particular genomic regions, we set a threshold for significant QTL hotspots based on the 99^th^ percentile of a Poisson distribution with a mean of 1.2 QTL (total QTL/total bins).

### Generation of near-isogenic lines (NIL)

NILs were generated by crossing selected RIAILs to each parental genotype. For each NIL, eight crosses were performed followed by six generations of propagating isogenic lines to ensure homozygosity of the genome. For each cross, PCR amplicons for insertion-deletion (indel) variants on the left and right of the introgressed region were used to confirm progeny genotypes and select non-recombinants within the introgressed region. NILs were whole-genome sequenced as described below to confirm their genotype (**File S7**). Reagents used to generate NILs and a summary of each introgressed region are detailed in the **Supplementary Information**. A statistical power calculation was used to determine the minimal number of technical replicates required to observe the predicted phenotypic effect of each QTL at 80% power. These calculations are listed in **Table S3**. The number of technical replicates tested per assay for any given toxin did not exceed 100 because of experimental timing constraints. The principal components that mapped to each NIL region are those with a QTL with a confidence interval that overlaps with or spans the entire introgressed region in the NILs (**Table 1, Table S4**).

**Table 1:** Toxins and principal components mapped per hotspot

| Toxin | Class | PCs in IVL | PCs in IVR | PCs in V |
| --- | --- | --- | --- | --- |
| Cadmium | Heavy Metal | 0 | 0 | 0 |
| Carmustine* | Chemotherapeutic | 1* | 0 | 1* |
| Chlorothalonil* | Pesticide | 2* | 1* | 1* |
| Chlorpyrifos | Pesticide | 1 | 1 | 0 |
| Cisplatin* | Chemotherapeutic | 2* | 1 | 2* |
| Copper | Heavy Metal | 2 | 0 | 0 |
| Diquat | Pesticide | 0 | 0 | 0 |
| Fluoxetine* | Neuropharmaceutical | 1 | 2* | 0 |
| FUdR | Chemotherapeutic | 1 | 1 | 0 |
| Irinotecan* | Chemotherapeutic | 0 | 1* | 2 |
| Mechlorethamine | Chemotherapeutic | 0 | 0 | 1 |
| Paraquat* | Pesticide | 0 | 0 | 1* |
| Silver* | Heavy Metal | 3* | 0 | 1* |
| Topotecan | Chemotherapeutic | 1 | 0 | 0 |
| Tunicamycin | Chemotherapeutic | 2* | 0 | 0 |
| Vincristine | Chemotherapeutic | 2 | 1 | 0 |

\*Denotes a toxin tested with NIL and/or CSS assays

### Whole-genome sequence library prep and analysis

DNA was isolated from 100-300 µL of packed animals using Qiagen’s Blood and Tissue kit (catalog # 69506). Following the ATL lysis step, 4 µl of 100 mg/mL RNAse was added to each sample and allowed to incubate for two minutes at room temperature. DNA concentration was determined using the Qubit dsDNA BR Assay Kit (catalog # Q32850). For each strain, a total of 0.75 ng of DNA was combined with 2.5 µL transposome (Illumina; kit # FC-121-1011) diluted 35x with 1x Tris Buffer (10x Tris Buffer: 100 mM Tris-HCl pH 8.0, 50 mM MgCl_2_) in a 10 µL final volume on ice. This reaction was incubated at 55°C for 10 minutes. The amplification reaction for each strain contained (final concentrations): 1x ExTaq Buffer, 0.2 mM dNTPs, 1 U ExTaq (Takara, catalog # RR001A), 0.2 µM primer 1, 0.2 µM primer 2, and 5 µL of tagmentation material from the previous step in a 25 µL total volume. Each strain had a unique pair of indexed primers. We first made a master mix containing buffer, water, dNTPs, and ExTaq then aliquoted the appropriate volume of this mix into each well. We added the specific primer sets to each well and finally the tagmentation reaction. The amplification reaction was incubated in a thermocycler with the following conditions: 72°C for three minutes (1x); 95°C for 30 seconds (1x); 95°C for 10 seconds, 62°C for 30 seconds, 72°C for three minutes (20x); 10°C on hold. We combined 8 µL from each amplification reaction to generate a pool of libraries. A portion of the libraries was electrophoresed on a 2% agarose gel. DNA was excised and gel purified using Qiagen’s Gel Purification Kit (catalog # 28706). The libraries were sequenced on the Illumina HiSeq 2500 platform using a paired-end 100 bp reaction lane. Alignment, variant calling, and filtering were performed as described previously (Cook, *et al.* 2016a). NIL and CSS genotypes were called using the VCF file and a Hidden Markov Model as described previously (Cook and Andersen 2017).

### Generation of chromosome substitution strains (CSS)

CSSs were generated by crossing N2 and CB4856 parental strains and mating cross progeny to each parental genotype. For each CSS, eight crosses were performed followed by six generations of propagating isogenic lines to ensure homozygosity of the genome. For each cross, PCR amplicons for indels on the left and right of the introgressed region were used to confirm progeny genotypes and select non-recombinants within the introgressed region. CSSs were whole-genome sequenced as described above to confirm their genotype (**File S7**). Reagents used to generate CSSs are detailed in the **Supplementary Information**. As described for NIL assays, power calculations were performed to determine the number of technical replicates required to observe the predicted phenotypic effect of the CSSs.

### Selection of traits to categorize in CSS and NIL assays

Pairwise correlations of RIAIL phenotypes among the 24 growth-response traits measured by the BIOSORT were calculated using the *cor* function within the *stats* package in R with the *use* argument set to “pairwise.complete.obs”. For each toxin, hierarchical clustering was performed using the function *hclust* from the *stats* package (R Core Team 2017). *Cutree* was then used to group the resulting dendrogram into *k* groups, where *k* is equal to the minimum number of principal components that explained at least 90% of the phenotypic variance in the RIAILs. For each principal component that mapped to a hotspot, the growth-response trait that was most correlated to that principal component, as well as all growth-response traits within that cluster of the dendrogram, were assayed in NIL and CSS experiments (**File S8, Table 2**).

**Table 2:** All traits tested in NIL and CSS assays

| PC | Hotspot | Correlated Traits | Correlation Range |
| --- | --- | --- | --- |
| carmustine.PC1 | V | mean.EXT, mean.TOF, q75.EXT, median.EXT, median.TOF, q75.TOF, median.norm.EXT, q90.TOF, q90.EXT | 0.72-0.95 |
| carmustine.PC6 | IVL | q25.norm.EXT, q10.norm.EXT | 0.33-0.39 |
| chlorothalonil.PC1 | V | mean.EXT, q75.EXT, mean.TOF, median.EXT, median.TOF, q75.TOF | 0.73-0.95 |
| chlorothalonil.PC2 | IVL | cv.TOF, cv.EXT | 0.72-0.90 |
| chlorothalonil.PC3 | IVL, IVR | mean.norm.EXT, q75.norm.EXT, q90.norm.EXT, median.norm.EXT | 0.50-0.65 |
| cisplatin.PC1 | IVL, V | mean.EXT, mean.TOF, median.EXT, median.TOF, q75.TOF, q75.EXT, q90.EXT, q90.TOF | 0.78-0.97 |
| cisplatin.PC3 | IVL | var.TOF, var.EXT | 0.38-0.54 |
| cisplatin.PC4 | V | norm.n, n | 0.76-0.80 |
| fluoxetine.PC1 | IVR | mean.norm.EXT, q75.norm.EXT, mean.EXT, q75.EXT, q90.norm.EXT, q90.EXT | 0.79-0.96 |
| fluoxetine.PC5 | IVR | q90.norm.EXT, q75.norm.EXT, mean.norm.EXT, q75.EXT, mean.EXT, q90.EXT | 0.07-0.40 |
| irinotecan.PC2 | IVR | cv.TOF, cv.EXT | 0.57-0.84 |
| paraquat.PC1 | V | median.EXT, mean.EXT, q25.EXT, q75.EXT, mean.TOF, q75.TOF, q10.EXT, q90.EXT, q90.TOF, median.TOF, q25.TOF, q10.TOF | 0.75-0.95 |
| silver.PC1 | V | mean.EXT, median.EXT, q75.EXT, mean.TOF, q90.EXT, q90.TOF, median.TOF, q75.TOF | 0.77-0.96 |
| silver.PC3 | IVL | q10.norm.EXT, q25.norm.EXT, mean.norm.EXT, median.norm.EXT, q75.norm.EXT, q90.norm.EXT | 0.32-0.64 |
| silver.PC4 | IVL | n, norm.n | 0.84-0.84 |
| silver.PC5 | IVL | n, norm.n | 0.41-0.41 |
| tunicamycin.PC1 | IVL | median.EXT, q75.EXT, mean.TOF, q75.TOF, median.TOF, median.norm.EXT, q90.EXT, q90.TOF, mean.EXT, q75.norm.EXT, mean.norm.EXT, q25.norm.EXT, q90.norm.EXT, q10.norm.EXT | 0.69-0.96 |
| tunicamycin.PC3 | IVL | norm.n, n | 0.47-0.50 |

### Categorization of CSS and NIL results

Toxin responses for NILs and CSSs were tested using the high-throughput fitness assay for traits correlated with mapped principal components as described above (**Table 2, File S9**). Complete pairwise statistical analyses of strains was performed for each trait tested in all CSS and NIL assays (Tukey honest significant difference (HSD) test, **File S10**). A *p-value* of *p* < 0.05 was used as a threshold for statistical significance. NIL recapitulation was defined by the significance and direction of effect of the NIL compared to the parental strains. Six categories were defined: 1) no parental difference, 2) recapitulation, 3) no QTL effect, 4) bidirectional interaction, unidirectional interaction, and 6) miscellaneous (**Table 3**). Traits for which N2 and CB4856 phenotypes were not statistically different comprise the ‘no parental difference’ category and were not further categorized. Traits in the ‘recapitulation’ category must satisfy the following criteria: significant difference between the parental strain phenotypes, significant difference between phenotypes of each NIL and the parent that shares its background genotype, and both NILs must display the expected direction of effect of the introgressed genotype. Traits with ‘no QTL effect’ displayed a significant parental phenotypic difference and the phenotype of each NIL was not statistically different from the phenotype of the parent sharing its background genotype. Traits that have a ‘bidirectional interaction’ must display a significant parental phenotypic difference, the phenotypes of both NILs must be significantly different from phenotypes of both parents, and the phenotypes of both NILs must be transgressive (lie beyond the phenotypic range of the parental strains). Lastly, traits with a ‘unidirectional interaction’ were categorized similarly to the bidirectional interaction, except only one NIL must display a transgressive phenotype and the other NIL either shows no QTL effect or recapitulation. Traits that did not fit these descriptions were categorized as ‘miscellaneous’.

**Table 3:** Categorization summary from NIL phenotypes

| Primary Category | Number of tests (99) |
| --- | --- |
| No Parental Effect | 23 |
| Recapitulation | 4 |
| No QTL Effect | 11 |
| Unidirectional Transgressive | 38 |
| Bidirectional Transgressive | 7 |
| Miscellaneous | 16 |

Traits in the chromosome V hotspot were further categorized using the combined data from both the CSS and NIL assays. Seven categories were defined: 1) no parental difference, 2) recapitulation, 3) no QTL effect, 4) external inter-chromosomal interaction (uni- or bidirectional), 5) internal inter-chromosomal interaction (uni- or bidirectional), 6) intra-chromosomal interaction (uni- or bidirectional), and 7) miscellaneous (**Table 4**). ‘No parental difference’ was defined by traits in which the parental strains were either not significantly different from each other or did not have the same direction of effect in both the CSS and NIL assays. ‘Recapitulation’ and ‘no QTL effect’ traits were defined by traits that were classified as either recapitulating or no QTL effect, respectively, in both assays. Traits displaying an ‘external inter-chromosomal interaction’ show evidence for interaction in the CSS but no interaction (either recapitulating or no QTL effect) in the NIL. On the other hand, traits displaying an ‘internal inter-chromosomal interaction’ showed evidence of the same interaction for both the CSS and the NIL assays. Finally, traits displaying an ‘intra-chromosomal interaction’ showed evidence of an interaction in the NIL but not in the CSS assay. All other traits that did not fit these descriptions were categorized as ‘miscellaneous’ (**File S11**).

**Table 4:**
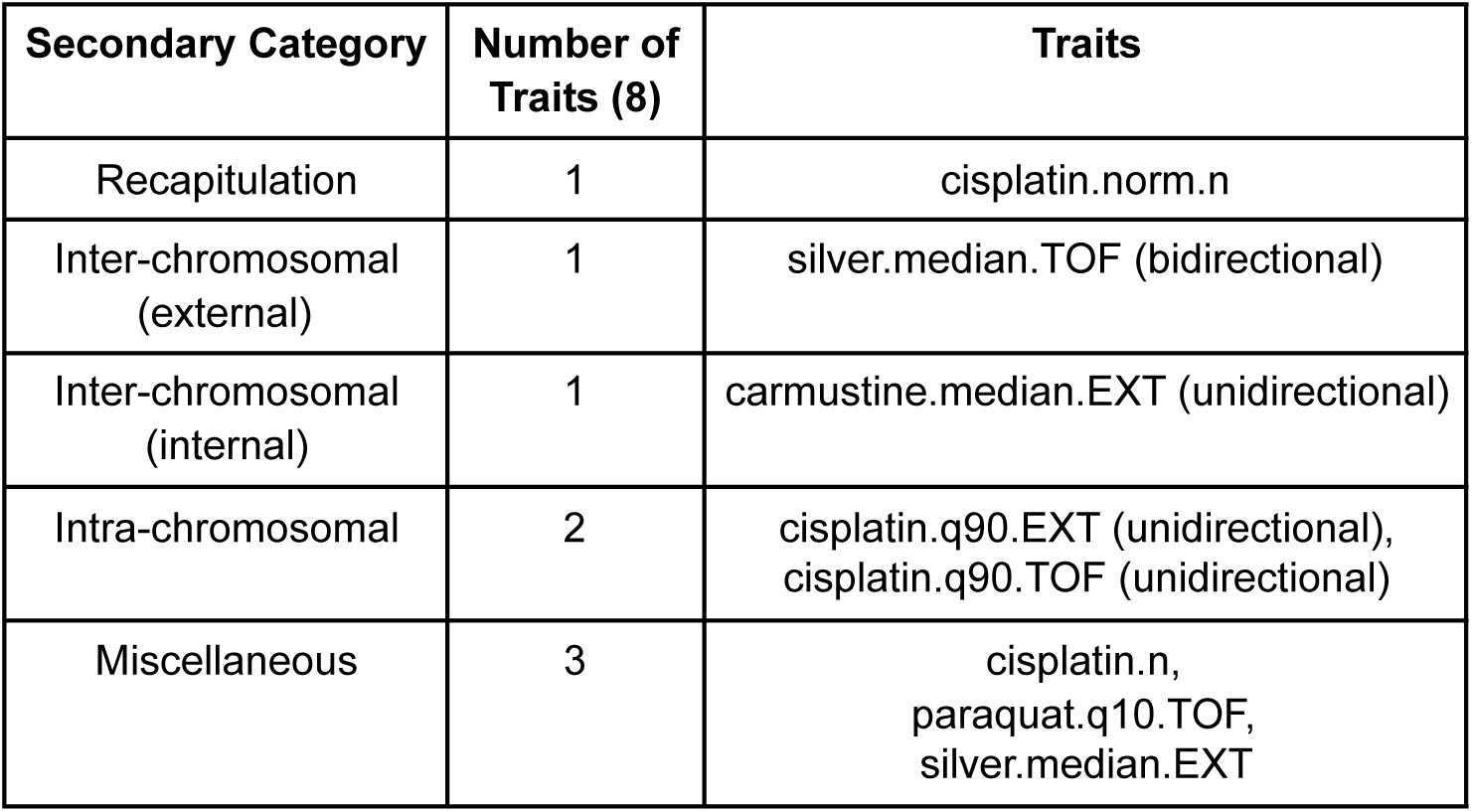
Categorization summary from combined NIL and CSS phenotypes

### Statistical analysis

All statistical tests of phenotypic differences in the NIL and CSS assays were performed in R (version 3.3.1) using the *TukeyHSD* function (R Core Team 2017) on an ANOVA model with the formula (*phenotype ∼ strain*). The *p-values* of individual pairwise strain comparisons were reported, and a *p-value* of *p* < 0.05 was deemed significant. The direction of effect of each NIL was determined by comparing the median phenotypic value of the NIL replicates to that of each parental strain. NILs whose phenotypes were significantly different from both parents and whose median lied outside of the range of the parental phenotype medians were considered hypersensitive or hyper-resistant. Comparing LOD scores and variance explained between traits with no parental effect and traits with a significant parental effect in the NIL assays was performed using a Wilcoxon rank sum test with continuity correction using the *wilcox.test()* function in R (R Core Team 2017).

### Data availability

**File S1** contains results of the dose response assays for all toxins. **File S2** contains the residual phenotypic values for each RIAIL for each trait. **File S3** contains the phenotypic values for each RIAIL for each of the significant principal components. **File S4** contains the annotated QTL and confidence intervals identified through linkage mapping. **File S5** contains the results of a two-factor genome scan for all traits with a significant QTL identified with linkage mapping. **File S6** contains the broad-sense heritability estimates as well as additive and interactive components of heritability for each trait. **File S7** is a VCF file for all NILs and CSSs mentioned in this manuscript. **File S8** contains each of the 97 significant PCs and the corresponding correlation value with each growth-response trait. **File S9** contains the residual phenotypic data for all strains, including parents, tested in the NIL and CSS assays. **File S10** contains the statistical significance for all pairwise combinations of strains tested for each trait. **File S11** contains the assay categorization for all traits tested with the NIL and CSS strains. The datasets and code for generating figures can be found at http://github/AndersenLab/QTLhotspot. All supplemental files, tables, and figures were uploaded to gsajournals.figshare.com.

## RESULTS

### Identification of QTL underlying variation in responses to 16 diverse toxins

Using a high-throughput fitness assay (Materials and Methods), we tested variation in 24 fitness-related traits in responses of four genetically divergent strains to different concentrations of 16 toxins, comprising chemotherapeutics, heavy metals, pesticides, and neuropharmaceuticals (**Figure S1, File S1, Table S1**). A concentration of each toxin was selected that minimized within-strain variation and maximized variation between two of these divergent strains, N2 (the laboratory strain) and CB4856 (a wild isolate from Hawaii) (**Table S 1**). For the selected concentration of each toxin, we assayed 24 growth-response traits for a panel of 296 recombinant inbred advanced intercross lines (RIAILs) generated between the N2 and CB 4856 parental genetic backgrounds (**File S2**) (Andersen et al. 2015). Because some of the growth-response traits are highly correlated (**Figure S2**), we performed principal component analysis (PCA) for each toxin. The minimum number of principal components (PCs) that explained at least 90% of the total phenotypic variance within each toxin was selected for mapping, for a total of 97 PCs across all toxins (minimum of five PCs and a maximum of eight PCs per toxin, **Table S2, File S3**). We then used linkage mapping to identify quantitative trait loci (QTL) that underlie variation in these 97 PCs.

We detected a total of 82 significant QTL (across 47 PCs) from the 97 PCs tested (**Figure 1, Figure S3, File S4**). We did not find a single toxin-response QTL shared robustly across all of the various PCs and toxins tested nor across all PCs within any one toxin. However, the majority of QTL on chromosome I were detected in responses to chemotherapeutics. Additionally, almost every toxin (with the exception of FUdR) had QTL that underlie trait variation on at least two different chromosomes, highlighting the diverse architectures implicated across traits, even within a single toxin. Despite the seemingly independent distributions of QTL, we found that the majority of the QTL (61%) mapped to chromosomes IV and V.

**Figure 1.**
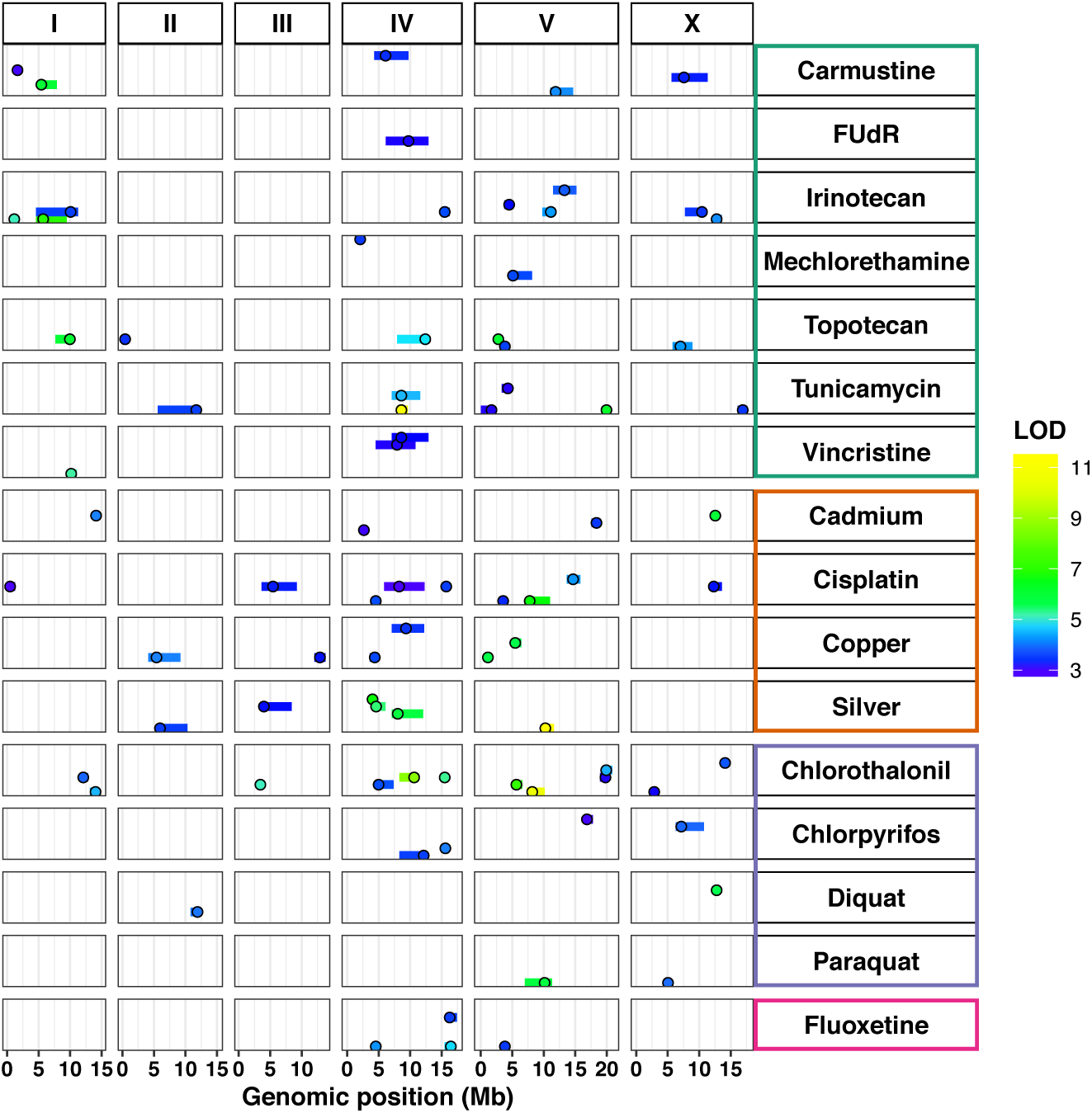
Diverse genetic architectures are implicated in responses to 16 toxins. Linkage mapping results for principal components that represent 82 QTL across 16 toxins, comprising chemotherapeutics (teal), heavy metals (orange), pesticides (purple), and neuropharmaceuticals (pink) are plotted. Genomic position (Mb) is shown along the x-axis, split by chromosome, and each of the 47 principal components with a significant QTL is plotted along the y-axis. Each QTL is plotted as a point at the location of the most significant genetic marker and a line indicating the 95% confidence interval. QTL are colored by the logarithm of the odds (LOD) score, increasing in significance from blue to green to yellow.

### Both additive and interactive QTL underlie toxin responses

For each of the PCs that were impacted by the 82 QTL identified using linkage mapping, we calculated the broad-sense heritability, the proportion of broad-sense heritability that could be attributed to additive genetic components (narrow-sense heritability) (**Figure 2A**), and the proportion of narrow-sense heritability that was explained by QTL detected through linkage mapping (**Figure 2B, File S6,** Materials and Methods). In many cases, additive genetic components could not explain all of the phenotypic variation predicted to be caused by genetic factors. These results suggest that other additive loci with small effect sizes impact toxin responses, but we failed to detect these QTL by our linkage mapping analyses, potentially because of high complexity and/or insufficient statistical power. Alternatively, this missing heritability could be indicative of genetic interactions (Bloom *et al.* 2013).

**Figure 2.**
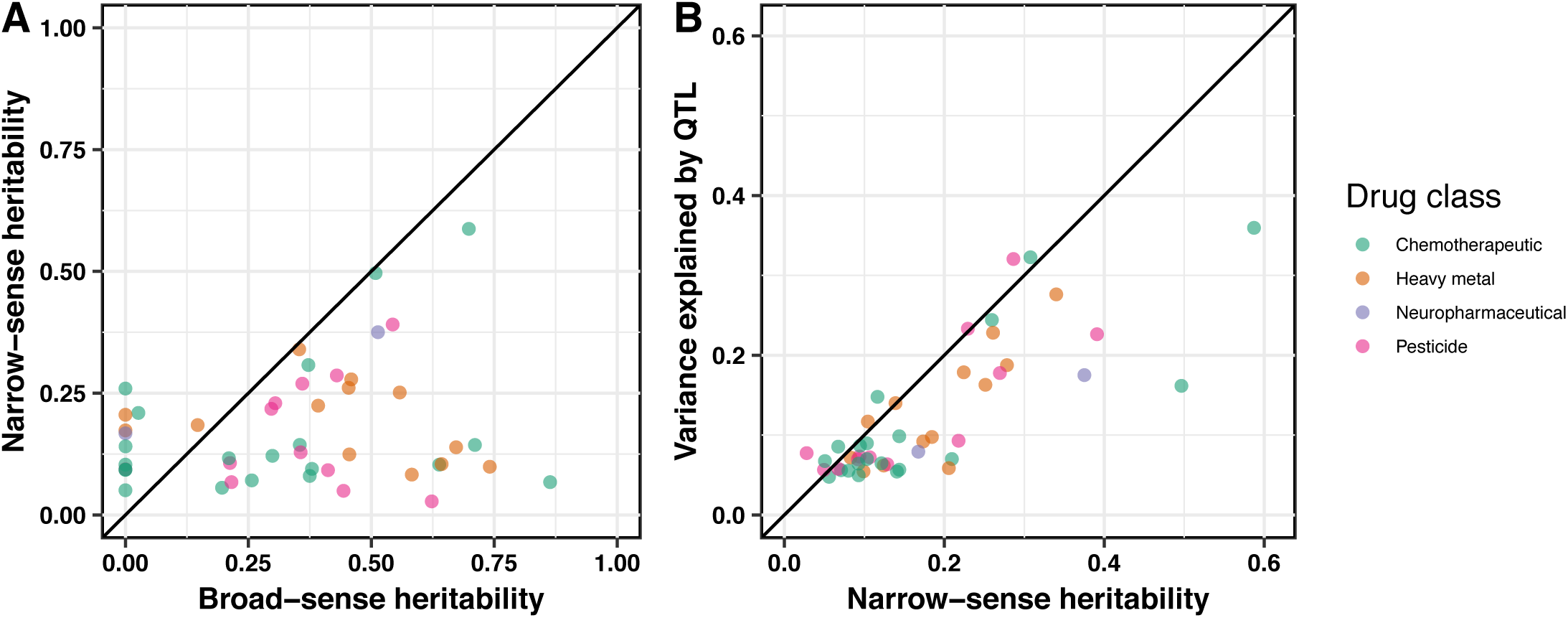
Additive genetic components identified by linkage mapping do not explain all heritable contributions to toxin-response variation. For 47 principal components representing the 82 QTL, we compared (**A**) the broad-sense heritability (x-axis) calculated from the RIAIL phenotypic data versus the narrow-sense heritability (y-axis) estimated by a mixed model and (**B**) the narrow-sense heritability (x-axis) versus the variance explained by all QTL detected by linkage mapping (y-axis). In both plots, each principal component is plotted as a point whose color indicates drug class (chemotherapeutic, heavy metal, neuropharmaceutical, or pesticide). The diagonal line represents y = x and is shown as a visual guide.

To determine how much of the phenotypic variance comes from additive or interacting genetic components, we fit a linear mixed-effect model to the RIAIL phenotype data for the 47 PCs controlled by the 82 QTL. We observed a range of additive and epistatic components contributing to phenotypic variation across toxin classes (**Figure S3, Figure S4, File S6**). On average, cisplatin, topotecan, and FUdR are primarily explained by additive models (**Figure S3**). Alternatively, paraquat, irinotecan, vincristine, and mechlorethamine have a larger fraction of their phenotypic variance attributable to genetic interactions than additive effects (**Figure S3**). To localize potential genetic interactions for these 82 QTL, we scanned the genome for interactions between pairs of markers that might affect the phenotypic distribution of the RIAIL panel (Materials and Methods). We identified three significant interactions (**File S5**). This two-factor genome scan was unable to localize all epistatic components identified by the linear mixed-effect model (**Figure 2**), perhaps because of missing small-effect additive loci in the model and/or insufficient statistical power to identify small-effect interactions.

### Three QTL hotspots underlie variation in responses to diverse toxins

The majority of toxin-response QTL cluster on chromosomes IV and V (**Figure 1**). We sought to determine if such QTL clustering could be expected by chance or if this clustering is indicative of toxin-response QTL hotspots. To account for the higher rate of recombination, and thus more genetic diversity, on the chromosome arms (Rockman and Kruglyak 2009), we divided the genome evenly into 65 bins and calculated the number of QTL that mapped to each bin (**Figure 3**, Materials and Methods). Three bins with more QTL than expected based on a Poisson distribution (Brem and Kruglyak 2005) were classified as hotspots. These hotspots are located on the center of chromosome IV, the right of chromosome IV, and the center of chromosome V and are hereby denoted as IVL, IVR, and V, respectively. Importantly, these hotspots are not driven by multiple principal components within a single toxin. Instead, hotspots comprise multiple QTL across a variety of principal components and toxins. In fact, 14 of the 16 toxins tested have a principal component that maps to at least one of the three hotspots (**Table 1**). Of the 82 QTL, 18 mapped to IVL, 8 mapped to IVR, and 9 mapped to V. In total, 33 QTL map to a hotspot (note that two QTL have confidence intervals that span both hotspots on chromosome IV). We sought to experimentally validate the predicted additive and epistatic effects on toxin responses for QTL that mapped to the three hotspots.

**Figure 3.**
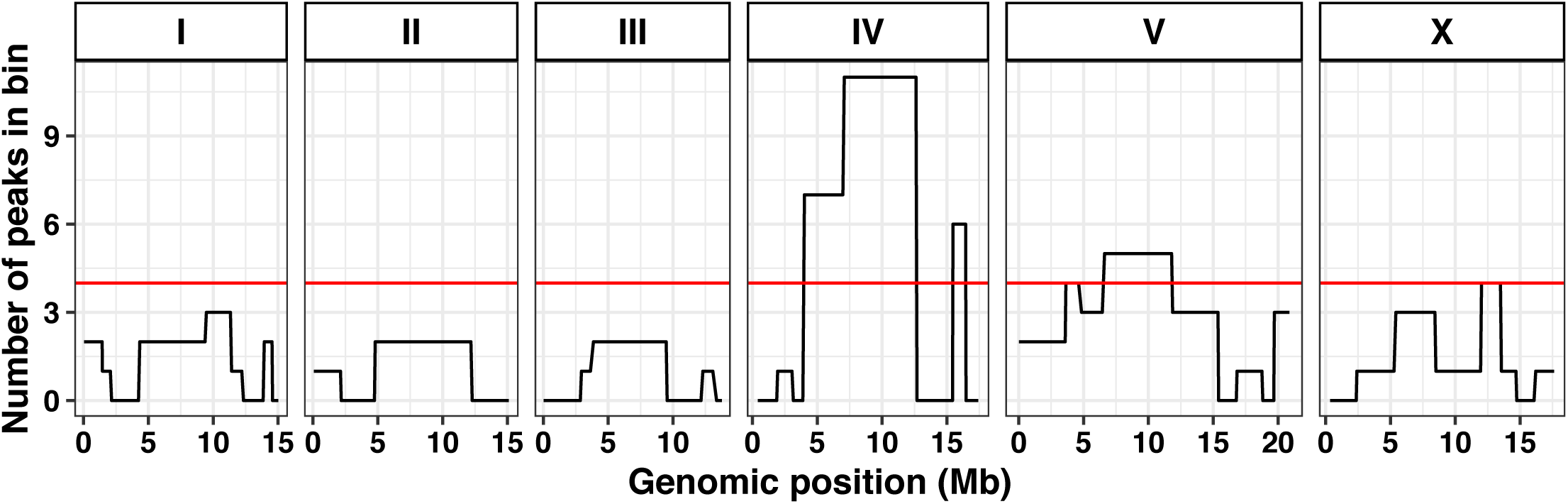
Three QTL hotspots impact toxin responses. Each chromosome is divided into equal bins of 26 cM, resulting in a total of 65 bins across the genome. The x-axis shows the genomic position (Mb), and the y-axis shows the number of QTL that lie within the corresponding bin. The red line indicates the 99^th^ percentile of a Poisson distribution with a mean of 1.26 QTL (total QTL/total bins).

### Near-isogenic lines recapitulate some of the predicted QTL effects

To experimentally validate the QTL identified from linkage mapping, we created near-isogenic lines (NILs) for the IVL, IVR, and V hotspots. Each NIL has a small genomic region introgressed from one parental strain into the genome of the opposite parental strain (Materials and Methods). These NILs were whole-genome sequenced and found to match the expected genotype in the hotspot region; however, occasionally additional breakpoints were observed (Material and Methods, **File S7**). We tested each NIL in our high-throughput fitness assay for a subset of the toxins with a QTL that maps to a given hotspot, choosing QTL with small, medium, and large effect sizes to test our ability to recapitulate various effect sizes (**Table 1, Table 2, Table S4**). We tested five toxins (ten QTL) with the IVL NILs, three toxins (four QTL) with the IVR NILs, and five toxins (six QTL) with the V NILs. In total, we tested 20 QTL across eight toxins for recapitulation using the NILs.

For each of these 20 QTL, we identified the toxin-response trait that is most correlated with the principal component controlled by that QTL. We then assayed the NILs for that toxin-response trait as well as all toxin-response traits within its same trait cluster, because each principal component comprises multiple toxin-response traits (**Table 2,** Materials and Methods). We tested 42 toxin-response traits with the IVL NILs, 12 toxin-response traits with the IVR NILs, and 45 toxin-response traits with the V NILs (**Figure S5, Table 2, File S9)**. In total, we performed 99 tests of recapitulation of QTL effects for toxin-response traits. The results of these tests allowed us to sort QTL effects into six different categories: ‘no parental effect’, ‘recapitulation’, ‘no QTL effect’, ‘unidirectional transgressive’, ‘bidirectional transgressive’, or ‘miscellaneous’ (**Figure S6**, **Table 3, File S11**).

Of these 99 tests, 23 did not display a significant phenotypic difference between the parent strains (N2 and CB4856) in the NIL assay and were categorized as ‘no parental effect’ (Materials and Methods, **Figure S5, Table 3**). The remaining 76 tests in which a significant parental difference was observed were classified further. We predicted that if a single QTL in the introgressed region contributed to the parental phenotypic difference, then each NIL would have a phenotype significantly different than the parental strain with the same genetic background. Furthermore, we expected each NIL to have a phenotype similar to the parental strain of its introgressed genomic region. This ‘recapitulation’ model was consistent for four tests (**Figure S5, Table 3**). The normalized brood size trait in cisplatin (cisplatin.norm.n in cisplatin PC4) is one such example of a trait in which the NILs on the center of chromosome V recapitulated the expected parental phenotype (**Figure 4A**). For 11 of the remaining 72 tests, the phenotype of each NIL was not significantly different from the phenotype of the parental strain sharing its background genotype (**Figure S5, Table 3**). This phenotype indicates that the introgressed NIL region was not affecting the toxin-response phenotype. This lack of QTL effect suggests that the genetic architecture is more complex, we lacked sufficient statistical power to detect the QTL effect, or the real QTL is outside the introgressed region. The NILs on the center of chromosome V showed this result for median animal length in silver (silver.median.TOF in silver PC1) (**Figure 4B**). The phenotypes of the NILs for the remaining 61 tests cannot be explained by a single QTL model. For many of these tests, we observed NIL phenotypes that are more sensitive or more resistant than both parental strains, suggesting that loci of opposite genotypes act additively or interact in the NILs to create transgressive phenotypes (Dittrich-Reed and Fitzpatrick 2013). This finding was supported by the mixed-effects model, which suggested that both additive and interacting QTL remained undetected by linkage mapping (**Figure 2**). We further explored the results of these 61 tests by characterizing them based on the patterns of the transgressive phenotypes we observed.

**Figure 4.**
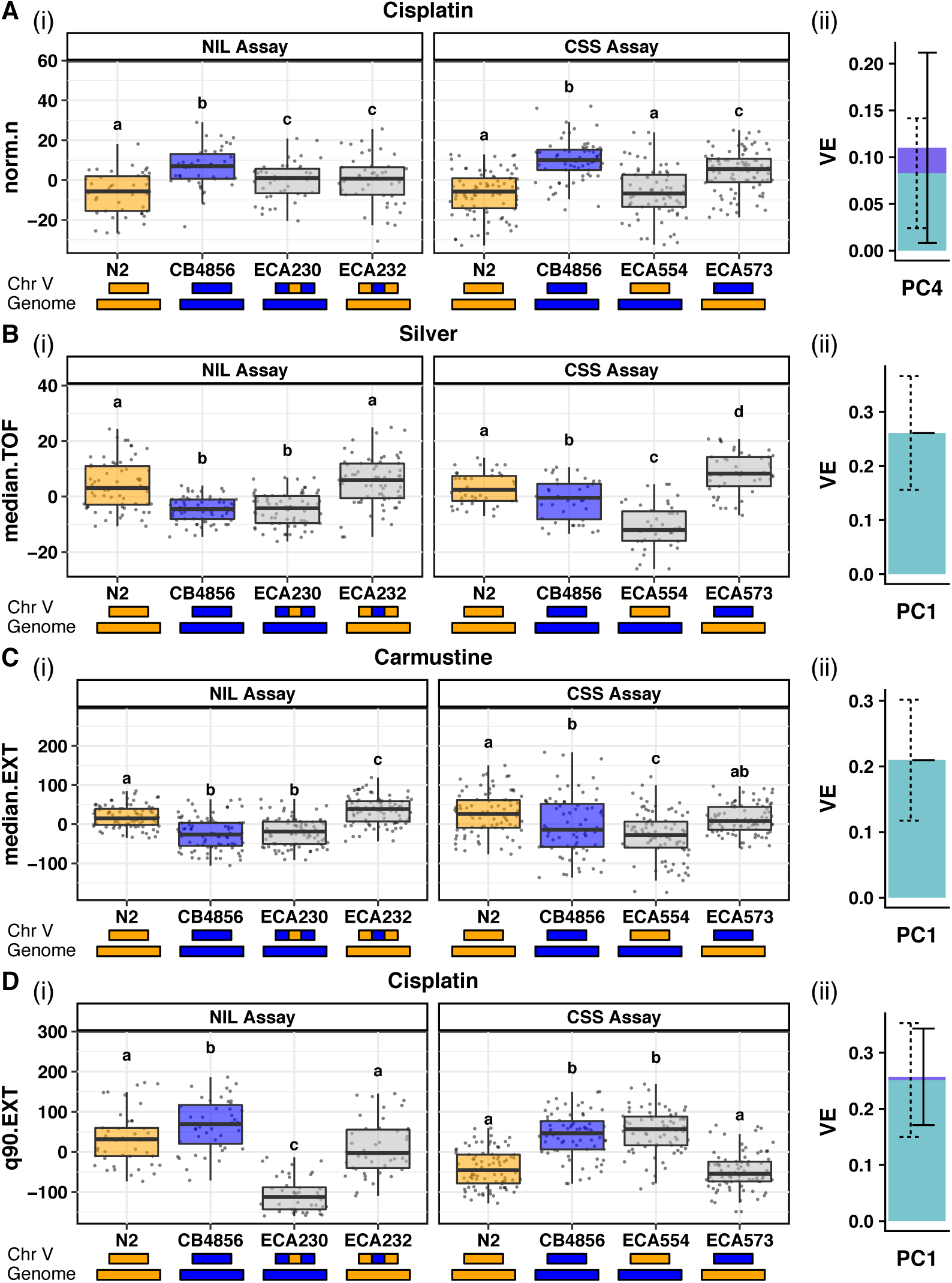
Results from near-isogenicline (NIL) and chromosome-substitution strain (CSS) tests of recapitulation of QTL effects are categorized based on potential genetic mechanisms implicated in toxin responses. A trait contributing to a mapped principal component for each category is reported: (**A**) Recapitulation (cisplatin norm.n, PC4), (**B**) Inter - chromosomal external bidirectionalloci (silver median.TOF, PC1), (**C**) Inter-chromosomal internal unidirectional loci (carmustine median.EXT, PC1), and (**D**) Intra - chromosomal unidirectional loci (cisplatin q90.EXT, PC1). In each case, we show results from (i) the NIL assay (left) and CSS assay (right) plotted as Tukey box plots. The y-axis indicates residual phenotypic values for the given trait. Different letters (a-d) above each Tukey box plot represent significant differences (p < 0.05) while the same letter represents non-significant differences between two strains (Tukey HSD). The genotype of each strain on the x-axis is modeled by the colored rectangles beneath the plots (N2 genotypes are orange, CB4856 genotypes are blue). (ii) A stacked bar plot shows the the proportion of pheno typic variation attributable to additive (light blue with dashed error bars) and interactive (dark blue with solid error bars) genetic factors of the principal component represented by each trait, based on a mixed model.

For 38 of these 61 tests, only one NIL showed a transgressive phenotype (**Table 3, Figure S5**). Some of these 38 ‘unidirectional transgressive’ phenotypes seem to show an antagonism that counteracted the effect of the introgressed region (a predicted sensitive phenotype becomes hyper-resistant or a predicted resistant phenotype becomes hypersensitive, *e.g.* carmustine.median.EXT in carmustine PC1, **Figure 4C**). Other phenotypes displayed synergy that increased the effect of the introgressed region (a predicted sensitive phenotype becomes a hypersensitive phenotype or a predicted resistant phenotype becomes a hyper-resistant phenotype, *e.g.* cisplatin.q90.EXT in cisplatin PC1, **Figure 4D**). Interestingly, in most cases (82%), the transgressive phenotype was observed in the strain with the N2 genotype introgressed into the CB4856 background.

In addition to unidirectional transgressive phenotypes, we identified seven tests with suggested ‘bidirectional transgressive’ phenotypes in which both NILs showed an extreme phenotype compared to the parental strains (**Figure S5**, **Table 3**). Some of these ‘bidirectional transgressive’ phenotypes were suggestive of purely antagonistic effects (*e.g.* tunicamycin.mean.norm.EXT, **Figure S5**), but others suggested an antagonistic effect in one NIL and a synergistic effect in the other (*e.g.* paraquat.median.TOF, **Figure S5**). We identified no cases of bidirectional synergistic effects. The remaining 16 tests of the 76 with a parental difference did not fall into any of the above categories and were classified as ‘miscellaneous’ (**Table 3**).

The toxin-response traits tested above for recapitulation of QTL effects were selected to represent principal components that were mapped with linkage mapping. We wanted to compare the NIL assay categorizations for the toxin-response traits that underlie each principal component to analyze the overall QTL effect (**Figure S7**). For example, two traits, n and norm.n, were selected to represent cisplatin PC4 (**Table 2**). Both of these toxin-response traits were placed into the ‘recapitulation’ category from the NIL assay results (**Figure S5, Figure S7**). These results suggest that a single additive QTL underlies the brood size variation captured by PC4. Fourteen tunicamycin-response traits were selected to represent tunicamycin PC1 (**Table 2**). Eight of these 14 traits displayed unidirectional transgressive phenotypes, four traits displayed bidirectional transgressive phenotypes, and the remaining two traits did not have a significant parental phenotypic difference (**Figure S5, Figure S7**). Regardless of the classification, we see the same trend of resistance (ECA231 > N2 > CB4856 > ECA229) across 11 of the 14 traits representing this principal component. Therefore, our strict significance thresholds for categorization might have caused some phenotypes to be mis-categorized (usually into the miscellaneous or no parental/QTL effect categories). The prevalence of transgressive phenotypes in tunicamycin-response traits suggests that multiple QTL, acting additively or interacting, might impact tunicamycin responses.

We next sought to compare categorizations of toxin-response traits and QTL effect sizes of the PCs for those traits. The QTL underlying cisplatin PC4 explains about 7% of the total phenotypic variance (**Table S4**). The traits selected to represent cisplatin PC4 were placed into the ‘recapitulation’ category, despite the small effect size of the QTL (**Figure S5, Figure S7**). In the other example above, the QTL underlying tunicamycin PC1 explains almost 16% of the total phenotypic variance, which is one of the highest effect sizes mapped in this study (**Table S4**). The toxin-response traits selected to represent this principal component showed mostly transgressive phenotypes, indicating undetected additive or interacting QTL despite the seemingly large-effect additive QTL identified in linkage mapping (**Figure S5, Figure S7**).

### Chromosome-substitution strains localize QTL underlying transgressive phenotypes

Because we found evidence of loci where opposite genotypes at each locus cause transgressive phenotypes, we attempted to further characterize these loci (**Figure 5, Figure S6**). To define each set of loci as either intra-chromosomal or inter-chromosomal, we built reciprocal chromosome-substitution strains (CSSs) for the hotspot on chromosome V that had the entire chromosome V introgressed from one parental strain into the genome of the opposite parental strain (Materials and Methods). The hotspot on chromosome V was chosen to isolate the effects of one hotspot and avoid complications arising from traits whose confidence intervals might lie within both of the hotspots on chromosome IV. The CSSs were whole-genome sequenced and found to have the expected genotype at all markers (Materials and Methods, **File S7**), except for the chromosome I incompatibility locus (Seidel *et al.* 2011; Seidel, Rockman, and Kruglyak 2008). We performed tests of recapitulation of QTL effects with the CSSs for each of the 45 toxin-response traits across the five toxins tested with the chromosome V NILs (**Figure S5, Table 2, Table S4, File S9**).

**Figure 5.**
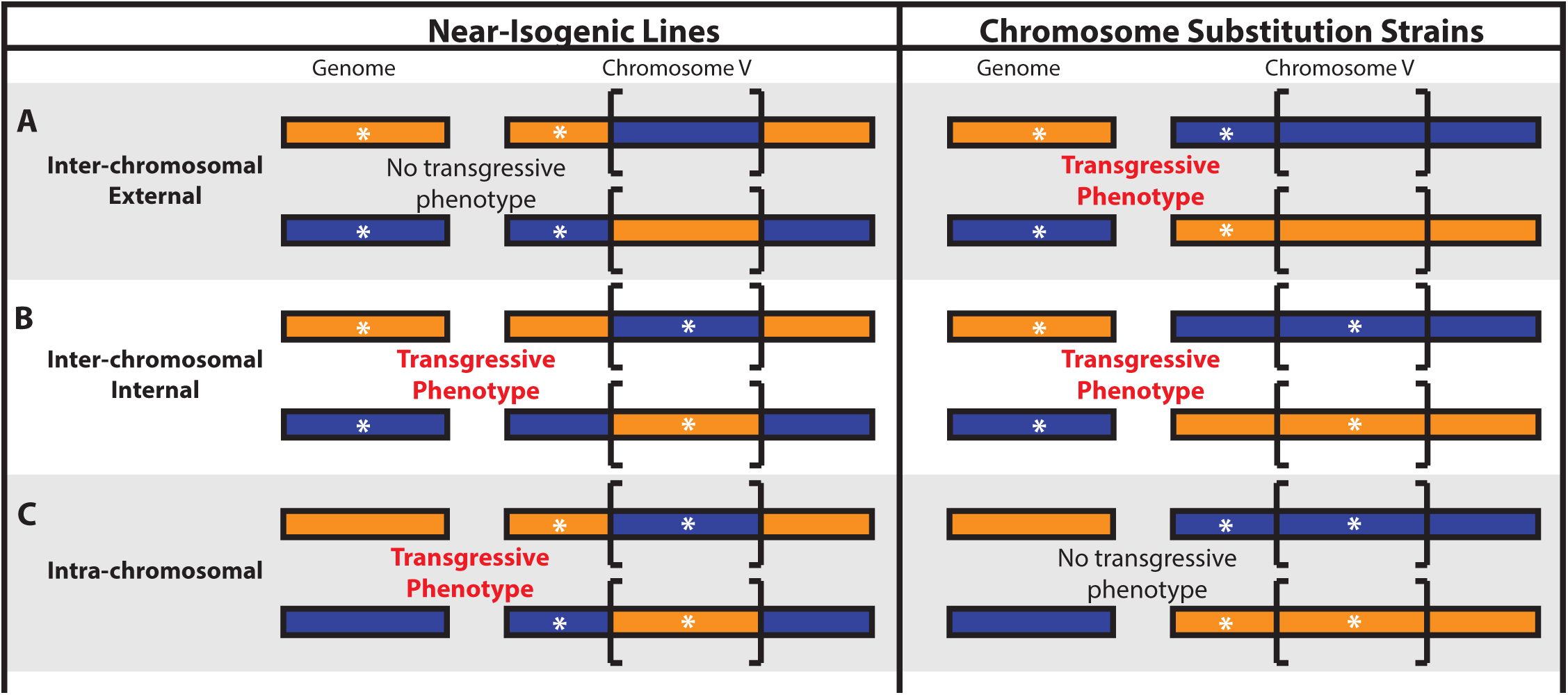
A model for potential locations of two loci is shown, according to toxin-response phenotypes of near-isogenic lines (NILs) and chromosome-substitution strains (CSSs). The NILs are represented on the left, and the CSSs are represented on the right. The strain genotype is indicated by colored rectangles. N2 is orange, and CB4856 is blue. Brackets indicate the genomic region that is introgressed in the NILs. White asterisks represent a potential location for additive or epistatic loci underlying transgressive phenotypes. Although bidirectional transgressive phenotype models are shown, each model could be bidirectional (both reciprocal introgressed strains show transgressive phenotypes) or unidirectional (only one reciprocal introgressed strain shows a transgressive phenotype). Models showing (**A**) inter-chromosomal external effects between a locus outside of the introgressed region in the NILs and a locus on another chromosome, (**B**) inter-chromosomal internal effects between a locus within the introgressed region in the NILs and a locus on another chromosome, and (**C**) intra-chromosomal effects between a locus within and a locus outside of the introgressed region in the NILs are drawn.

For traits in which the parental phenotypic difference was significant and consistent across the NIL and CSS tests, NIL and CSS phenotypes could be compared across assays. Eight traits across five toxins fit this criterion (**File S11, Table 4**). One trait (cisplatin.norm.n) displayed phenotypic ‘recapitulation’ of the introgressed region in both the NIL and the CSS tests, suggesting a single QTL model (**Figure 4A, Table 4**). Alternatively, transgressive phenotypes are indicative of a multi-QTL model, and locations of additive or interacting QTL can be surmised by comparing results from the NIL and CSS tests. Transgressive phenotypes controlled by inter-chromosomal loci are defined by two loci on separate chromosomes that act additively or epistatically. Because NILs and CSSs have introgressed genotypes on chromosome V, we can deduce that at least one of the two inter-chromosomal loci is located on chromosome V. We further divided the inter-chromosomal class into two categories: ‘inter-chromosomal external’, in which the chromosome V locus is outside the region introgressed in the NILs (**Figure 5A**) and ‘inter-chromosomal internal’, in which the chromosome V locus is within the region introgressed in the NILs (**Figure 5B**). For an ‘inter-chromosomal external’ model, we expect only the CSSs to display hypersensitivity or hyper-resistance, because both loci share the same genotype in the NILs (**Figure 5A**) and would therefore not result in a more extreme phenotype than both parents. We found one such trait that fits a ‘bidirectional inter-chromosomal external’ loci model (silver.median.TOF) (**Figure 4B, Table 4**). For an ‘inter-chromosomal internal’ model, we expected both the CSSs and the NILs to display the same hypersensitivity or hyper-resistance, because both strains share the same genotype across the introgressed region in the NILs (**Figure 5B**). We identified one such trait that fits a ‘unidirectional inter-chromosomal internal’ loci model (carmustine.median.EXT) (**Figure 4C, Table 4**). To identify intra-chromosomal loci that underlie transgressive phenotypes in the remaining 10 traits, we searched for traits that display evidence of either a uni- or bidirectional transgressive phenotype in the NILs but not in the CSSs (**Figure 5C**). This result would suggest that two loci of opposite genotypes on chromosome V, one within and one outside the region introgressed in the NILs, act additively or epistatically to cause transgressive phenotypes. We found two examples of such ‘unidirectional intra-chromosomal’ loci models (*e.g.* cisplatin.q90.EXT, **Figure 4D, Table 4**). The remaining three traits could not be characterized beyond their NIL assay characterization based on the results of the CSS assay (**Table 4**).

We revisited the two-factor genome scan results for each of these eight empirically classified traits and compared the findings from these two independent methods used to identify multiple additive or epistatic QTL. No traits with significant interaction terms were identified by the two-factor genome scan. Although many other pairs of loci show suggestive evidence of additive or interacting effects (**File S5**), an increase in statistical power is required to definitively compare these suggestive findings to our empirically derived model. Overall, this study highlights the benefits of leveraging both experimental and computational strategies to further dissect genetic components that underlie quantitative traits in a metazoan model.

## DISCUSSION

Here, we show that three QTL hotspots underlie differences in responses to 16 diverse toxins. We further characterized these QTL using both modeling and empirical approaches. Through the use of near-isogenic lines and chromosome-substitution strains, we confirmed small-effect QTL and attempted to identify and localize genomic regions causing transgressive phenotypes. Finally, we used statistical analyses to computationally identify loci that might support some of our empirical findings. Although the number of biological replicates and recombinant strains in this study increased our power to detect QTL compared to previous studies, we are still too underpowered to definitively assess if missing heritability is composed of small additive effects or genetic interactions.

### Pleiotropic regions underlie QTL shared between and among toxin classes

We performed principal component analysis on toxin-response phenotypes collected for a panel of RIAILs and used linkage mapping to identify 82 toxin-response QTL. Although some of these QTL are unique to one particular toxin, others suggest the existence of pleiotropic QTL that underlie responses to a diverse set of toxins. In particular, three QTL hotspots across chromosomes IV and V were enriched for toxin-response QTL and were investigated further. Because the molecular mechanisms implicated in responses to each toxin differ drastically, the notion that a single gene in each hotspot is regulating the response to several toxins is unlikely. However, the possibility exists that a single gene involved in drug transport could underlie one or several of these hotspots. More likely, multiple genes in close proximity, each regulating a process controlling cellular proliferation and survival, might underlie these hotspots. Notably, two of the three QTL hotspots are in swept regions with lower genetic diversity at the species level (Andersen *et al.* 2012; Laricchia *et al.* 2017; Cook, *et al.* 2016a; Cook, *et al.* 2016b). The laboratory strain, N2, has experienced each of the selective sweeps, and CB4856 has not. Linkage mapping using a panel of RIAILs built between these two strains could identify QTL that underlie phenotypic differences between swept and non-swept strains. Moreover, identifying QTL in these swept regions that underlie variation in fitness-related traits might indicate selective pressures that could have led to these chromosomal sweeps. For example, N2 is more resistant than CB4856 to tunicamycin (**Figure S5**), an antibiotic and chemotherapeutic produced by the soil bacterium *Streptomyces clavuligerus* (Price and Tsvetanova 2007). This result might suggest that selective pressure toward responses to antibiotic compounds played a role in driving resistance-conferring alleles, such as those present in N2, to a high frequency. Alternatively, climate conditions could also impact local niche environments to sensitize toxin responses (Evans *et al.* 2017). We observed that N2 is more resistant than CB4856 in responses to the majority of conditions, which could indicate that alleles present in swept strains confer robustness in responses to many conditions. This result emphasizes the importance of genetic background when considering toxin effects (Zdraljevic and Andersen 2017).

In addition to the three QTL hotspots, pleiotropic QTL across toxins within certain classes are suggested by our linkage mapping results. We observed an enrichment of QTL from the chemotherapeutic class on chromosome I, which could be representative of QTL that underlie a common mechanism targeted by these toxins, such as DNA damage or cell-cycle control. However, because many of these chemotherapeutics have distinct mechanisms of action and share these mechanisms with other toxin classes, this enrichment is likely caused by an overrepresentation of chemotherapeutics in our study. Direct comparisons of toxins with similar cellular mechanisms could provide more insights. For example, irinotecan and topotecan are both chemotherapeutics that cause DNA damage by inhibiting topoisomerase I (Pommier 2006) and share a QTL on the center of chromosome I. However, each of these chemotherapeutics also maps to distinct regions of the genome. For example, the irinotecan-response QTL on the right arm of chromosome V is not mapped for topotecan response and the topotecan-response QTL on the left arm of chromosome II is not mapped for irinotecan response. Vincristine also maps to this same region, however its mechanism of action is distinct from irinotecan and topotecan. The combination of overlapping and distinct genetic architectures underlying these highly similar compounds suggest that although some genetic variation implicated in responses to irinotecan and topotecan is shared, other QTL are specific to each compound and not representative of a general topoisomerase I inhibition mechanism. We have also observed this phenomenon of distinct genetic architectures underlying similar compounds for benzimidazole responses (Zamanian *et al.* 2018).

### A multi-faceted approach suggests that undetected epistatic loci impact toxin responses

To determine if we had sufficient power to experimentally validate even small-effect QTL, we constructed NILs for the three hotspots and assayed them in responses to multiple toxins. Because each principal component comprises multiple toxin-response traits, we measured NIL phenotypes for the most correlated toxin-response traits for each principal component to test recapitulation of QTL effects. For some of these tests of recapitulation for small-effect QTL, NILs showed a significant phenotypic effect. One such example is cisplatin.norm.n and cisplatin.n which represent the QTL mapped by cisplatin.PC4 that only explains 7% of the phenotypic variance. Our ability to recapitulate such a small effect suggests that our assay had sufficient power to detect small phenotypic effects in at least some cases. We postulated that our inability to recapitulate other QTL effects could be attributed to either insufficient power or additional additive or epistatic QTL that were undetected by linkage mapping. Particularly in cases where the NILs displayed transgressive phenotypes, undetected loci of opposite genotypes, acting additively or epistatically, likely caused these effects. Therefore, we investigated these interactions and found evidence for additional QTL that interact with the originally detected loci. However, we must note that whole-genome sequence data revealed that three of our NILs had a portion of the genome from the background of the starting RIAIL (**File S7**). Although we do not believe that these small regions are responsible for the unexpected phenotypes observed, this explanation could be a consideration for certain silver, cisplatin, carmustine, and chlorothalonil PCs, as they have significant QTL in these identified regions. This example emphasizes the importance of whole-genome sequencing NILs to verify the expected genotypes before making conclusions about phenotypic effects of a targeted QTL.

We used the results from the NIL assays to classify each test into a category that predicts a genetic model that might underlie NIL phenotypes. Categorizations were consistent across traits representing a principal component, with most of these traits falling into one or a few categorizations. This widespread consistency suggests that similar genetic architectures underlie phenotypes for these grouped traits. Furthermore, this consistency highlights the reproducibility of our high-throughput toxin response assay, because results from independent assays (trait correlations, linkage mappings from RIAIL assays, and phenotype classifications from NIL assays) often align to support the same conclusion obtained from the individual experiments.

The majority of cases of transgressive phenotypes occurs when the N2 genotype is introgressed into the CB4856 genome. This trend might indicate allele-specific unidirectional incompatibilities between the two strains, and localizing these interactions could improve our understanding of the evolutionary processes driving such incompatibilities. However, identifying the loci that underlie these unidirectional transgressive phenotypes using a mixed-effect model or a two-factor genomic scan is difficult, because only a small number of the RIAILs have the required allelic combinations to quantify such an effect. For example, cisplatin.q90.EXT, a trait chosen to represent cisplatin PC1, fits a unidirectional intra-chromosomal model. The results of the NIL and CSS assays show that, although the CSSs seem to display no QTL effect, the NIL with the N2 genotype introgressed into the CB4856 genome displays strong hypersensitivity (**Figure 4D**). All of the narrow-sense heritability for cisplatin PC1 (25%) predicted by the mixed-effect model is explained by the three QTL identified through linkage mapping (the variance explained estimates of these three QTL add up to 26%, **File S4, File S6**). This finding suggests that most of the additive loci have been identified through linkage mapping. Therefore, the intra-chromosomal loci are likely acting epistatically to cause a unidirectional transgressive phenotype. However, using our mixed-model approach, we do not find a significant interaction component for cisplatin PC1, the principal component that is represented by cisplatin.q90.EXT. A two-dimensional genome scan for multiple loci that underlie cisplatin PC1 provides suggestive evidence for a two-QTL model over a one-QTL model, with or without interaction between the loci (**File S5**). These two loci are located on the left of chromosome V (outside the NIL interval) and in the center of chromosome V (inside the NIL interval) and match our empirical evidence of two intra-chromosomal loci underlying the transgressive phenotype observed (**Figure 5C**). Because the transgressive phenotype is unidirectional, RIAILs without the allelic combination that causes extreme phenotypes could dilute our power to detect the loci. For this reason, combining both computational models and empirical investigation facilitates the detection of loci that control transgressive phenotypes. Additionally, future studies should include even larger RIAIL panels than what we used here to empower approaches to investigate the contributions of interactive loci.

Although we are statistically underpowered to identify some small-effect additive and interacting loci through modeling, the combination of three methods of searching for potential interactions suggests that not all fitness traits in *C. elegans* are composed of additive effects. Our two computational methods were used to identify additive and epistatic loci underlying many toxin responses, but their power was limited in cases of unidirectional transgressive phenotypes. Alternatively, the NIL and CSS phenotypic assays were able to identify unidirectional transgressive phenotypes, but they were restricted by their inability to distinguish between additive and epistatic loci. Constructing double CSS strains or multi-region NILs in which pairwise combinations of two genomic regions are introgressed within the opposite genotype could help to further define loci underlying transgressive phenotypes. However, each locus must be isolated to determine if the two loci act additively or epistatically. The results from the two-dimensional genome scan might provide insights into where to begin this approach. In cases where all three of our techniques suggested epistasis, we suspect that these QTL are not purely additive. Generating an even larger panel of recombinant strains and assaying a much larger number of biological replicates might allow us to further address the debate about how heritable loci contribute to trait variation in metazoans.

## ACKNOWLEDGEMENTS

The authors would like to thank Bryn Gaertner, Samuel Rosenberg, Tyler Shimko, and Robyn Tanny for assistance on mapping drug sensitivities and members of the Andersen lab for helpful comments on this manuscript. This work was supported by the following grants to E.C.A.: National Institutes of Health R01 subcontract to ECA (GM107227), the Chicago Biomedical Consortium with support from the Searle Funds at the Chicago Community Trust, and an American Cancer Society Research Scholar Award (127313-RSG-15-135-01-DD). S.C.B. was supported by the Biotechnology Training Grant (T32GM008449). K.S.E. was supported by the Cell and Molecular Basis of Disease training grant (T32GM008061). J.S.B. was supported by the Howard Hughes Medical Institute. The funders had no role in study design, data collection and analysis, decision to publish, or preparation of the manuscript.

